# Physical localization of 45S rDNA in *Cymbopogon* and the analysis of differential distribution of rDNA in homologous chromosomes of *Cymbopogon winterianus*

**DOI:** 10.1101/2021.08.25.457632

**Authors:** Shivangi Thakur, Upendra Kumar, Rashmi Malik, Darshana Bisht, Priyanka Balyan, Reyazul Rouf Mir, Sundip Kumar

**Affiliations:** Department of Molecular Biology and Genetic Engineering, College of Basic Sciences and Humanities, G. B. Pant University of Agriculture & Technology, Pantnagar-263145, India; Department of Molecular Biology, Biotechnology and Bioinformatics, CCS Haryana Agricultural University, Hisar-125004, India; Department of Genetics and Plant Breeding, College of Agriculture, G. B. Pant University of Agriculture & Technology, Pantnagar-263145, India; Department of Botany, Deva Nagri P.G. College, CCS University Meerut-245206, India; Division of Genetics & Plant Breeding, Sher-e-Kashmir University of Agricultural Sciences & Technology of Kashmir (SKUAST-Kashmir), Srinagar-190025, (J&K), India

## Abstract

*Cymbopogon*, commonly known as lemon grass, is one of the most important aromatic grasses having therapeutic and medicinal values. FISH signals on somatic chromosome spreads off *Cymbopogon* species indicated the localization of 45S rDNA on the terminal region of short arms of a chromosome pair. A considerable interspecific variation in the intensity of 45S rDNA hybridization signals was observed in the cultivars of *Cymbopogon winterianus* and *Cymbopogon flexuosus*. Furthermore, in all the varieties of *Cymbopogon winterianus* namely Bio-13, Manjari and Medini, a differential distribution of 45S rDNA was observed in a heterologous pair of chromosome 1. The development of *Cymbopogon winterianus* var. Manjari through gamma radiation may be responsible for breakage of fragile rDNA site from one of the chromosomes of this heterologous chromosome pair. While, in other two varieties of *Cymbopogon winterianus* (Bio-13 and Medini), this variability may be because of evolutionary speciation due to natural cross among two species of *Cymbopogon* which was fixed through clonal propagation. However, in both the situations these changes were fixed by vegetative method of propagation which is general mode of reproduction in the case of *Cymbopogon winterianus*.

## Introduction

*Cymbopogon*, commonly known as lemon grass, is one of the most important aromatic grasses belonging to family *Poaceae* with proven therapeutic and medicinal values. It is grown for commercial and industrial purposes in tropics and subtropics of Asia, America and Africa [1]. Industrial interest in citronella oils is due to extensive use of its various components as fragrance in perfumes, soaps, mosquito repellent and as flavour additives in food products. *Cymbopogon* species display wide variation in morphological attributes and essential oil composition at inter and intraspecific levels [2]. The first cytological study in Indian *Cymbopogon* to ascertain the chromosome number [3-4] indicated the different ploidy levels in these genera varying from diploid (2n=20) to tetraploid (2n=40) and hexaploid (2n=60). However, as per available reports in literature, the cytogenetic studies in this genus have been limited to chromosome count and preliminary karyotype description of the cultivated species. The characterization of *Cymbopogon* germplasm largely been done on phenotypic characteristics [5] and more recently on the basis of some molecular markers such as RAPD and SSR [6]. Analysing the genome organization of plants reveal evolutionary relationships of different genomes, which may also be useful for crop improvement. The ribosomal RNA genes represent two highly conserved tandemly arrayed gene families namely 45S rDNA and 5S rDNA which have been studied extensively in plant genomes. Because of the numerous copies of these highly conserved families of repeated sequences, their physical location on the chromosome can be easily visualized. The 45S rDNA sites in somatic chromosomes are most extensively utilized and widely documented chromosomal regions in eukaryotes through fluorescent *in situ* hybridization (FISH). The 45S rDNA family together with the intergenic spacer (IGS) is present as tandem arrays within the nucleolus organizer regions (NORs) of satellite chromosomes and also at other chromosomal sites where they may not be associated with NOR [7-9]. Length polymorphism of these repeat units has been reported in a wide range of plants and animals and are attributed to variation in number of sub-repeats that are found in IGS. The length of IGS and the chromosomal location of 45S rDNA genes are often characteristic of a species and were widely used to study the phylogenetic relationships of several plant species [10-13]. In view of the limited cytological reports on *Cymbopogon*, molecular cytogenetic studies for the identification of individual chromosomes are urgently needed in this important crop. Therefore, the present investigation was conducted using 45S rDNA as a probe to develop valuable FISH landmarks of somatic chromosomes of *Cymbopogon* which may be utilized in subsequent molecular cytogenetic studies to generate physical maps of *Cymbopogon* species.

## Results

### Karyotype analysis

All the four varieties studied during the present investigation were observed as diploid with chromosome number (2*n*=20). Variety Bio-13 belonging to *C. winterianus* is one of the supreme significantly commercialized variety of Java Citronella grass. This variety of *C. winterianus* contain diploid chromosome complement as 2*n=*20 with basic chromosome number *x* = 10 (Fig. 1a, 1c and 2a). Arm ratios for the chromosomes of this species ranged from 1.05 to 3.37 (Fig. 2a) and the range of chromosome lengths lies between 1.3 to 3.12 µm. Bio-13 observed to have 3 chromosomes in the range of 2.0 to 3.0 µm (3C) and rest of the seven chromosomes in the range of 1 to 2 µm (7D). As per the arm ratios of different chromosomes of this variety, 3 chromosomes were metacentric and 7 were sub-metacentric (Table 1).

**Figure 1.**
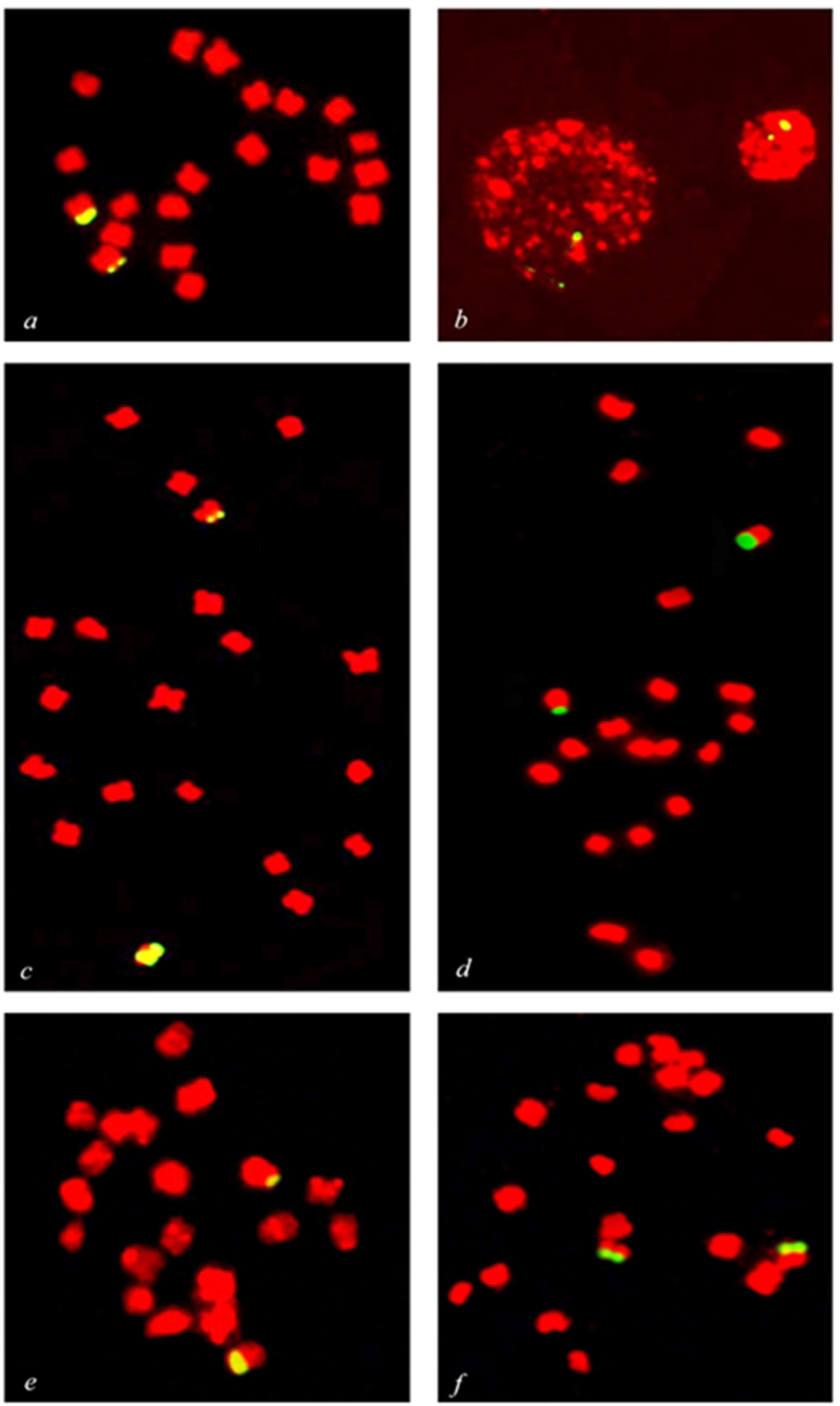
Mitotic cells of variety Bio-13, Medini, Manjari (*C. winterianus*) and Krishna (*C. flexuosus*) after fluorescence *in situ* hybridization with 45S rDNA (yellow/green) as FISH probe. The chromosomes were counterstained with propidium iodide (red). **(a and c)** Somatic metaphase showing diploid chromosomes of Bio-13 (2*n =* 20) showing differential hybridization signals of 45S rDNA on a pair of somatic chromosomes. **(b)** Interphase nuclei of Bio-13 showing differential hybridization signals of 45S rDNA **(d)** Somatic metaphase showing diploid chromosomes of Manjari (2*n =* 20) showing differential hybridization signals of 45S rDNA on a pair of somatic chromosomes.**(e)** Somatic metaphase showing diploid chromosomes of Medini (2*n =* 20) showing differential hybridization signals of 45S rDNA on a pair of somatic chromosomes. **(f)** Somatic metaphase showing diploid chromosomes of Krishna (2*n=*20) showing hybridization signals of 45S rDNA on a pair of somatic chromosomes.

**Figure 2.**
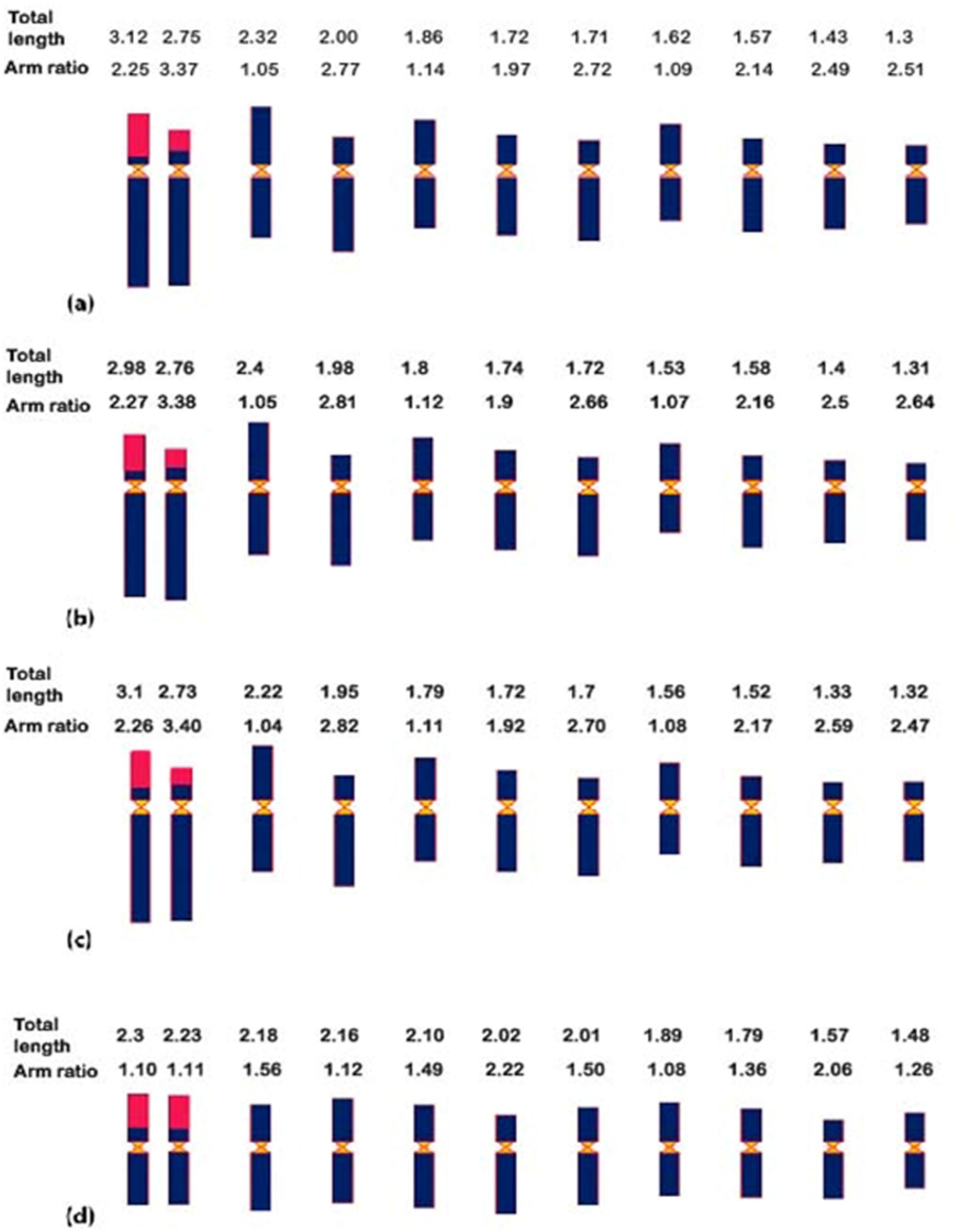
Total length and arm ratio of chromosome. Schematic representation of chromosome arranged in descending order of their length: **a**. Karyogram of Bio-13 (*C. winterianus*), **b**. Karyogram of Manjari (*C. winterianus*), **c**. Karyogram of Medini (*C. winterianus*), **d**. Karyogram of Krishna (*C. flexuosus*).

**Table 1.**
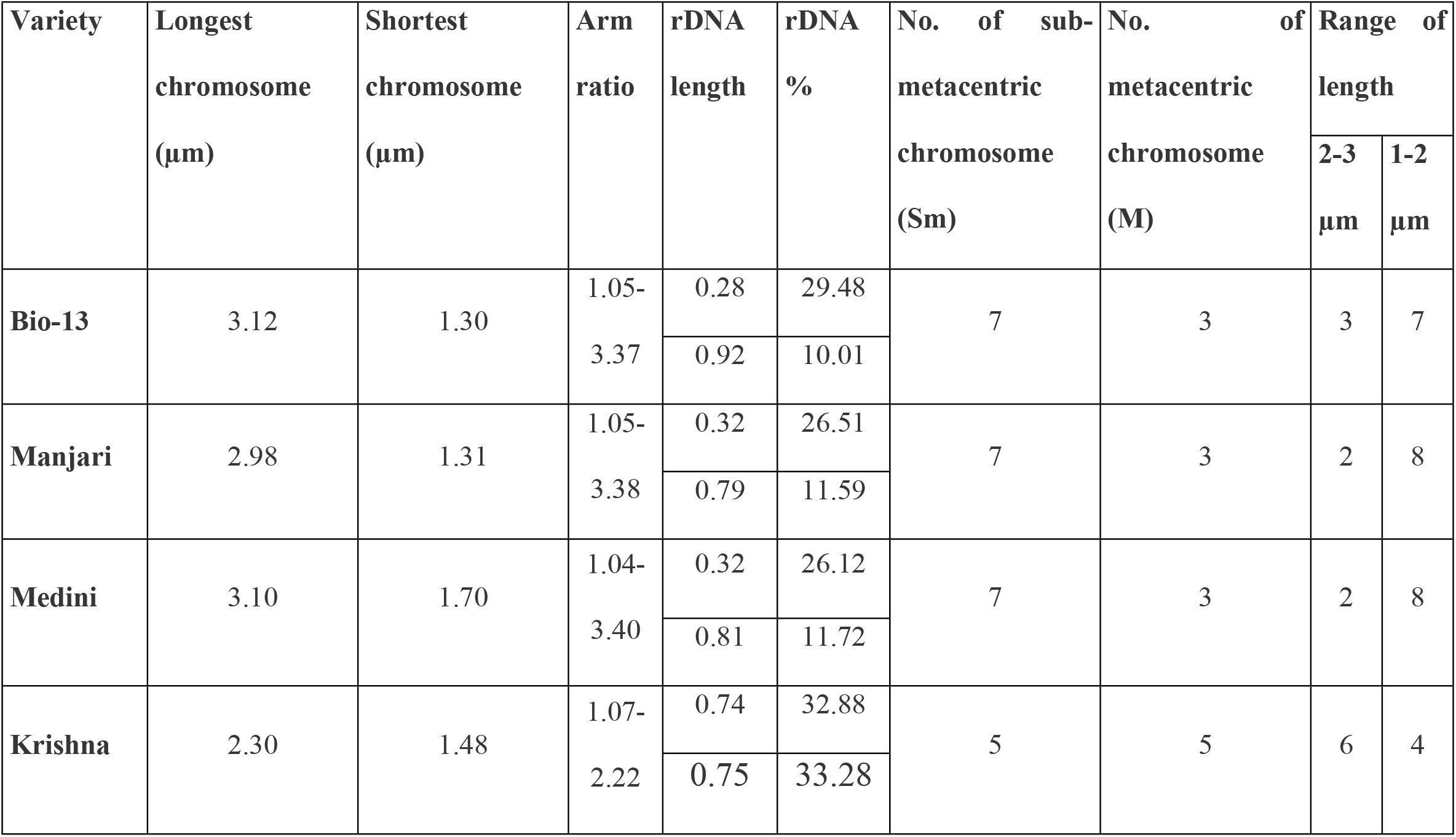
Description of karyotyping in all the varieties.

Manjari (*C. winterianus*) also shows its somatic chromosome complement as 2*n*=20 (diploid) with basic chromosome number *x*=10 (Fig. 1d). Arm ratios for the chromosomes of this species ranged from 1.05 to 3.38 µm and the range of chromosome lengths lies between 1.31 to 2.98 µm (Fig. 2b) which is almost equivalent to *C. winterianus* var. Bio-13. This variety is observed to have 2 chromosomes in the range of 2.0 to 3.0 µm (2 C chromosomes) and rest of the eight chromosomes ranged in 1.0 to 2.0 µm (8 D chromosomes). As per the arm ratios (Table 1) of different chromosomes of this variety, 3 chromosomes were metacentric and 7 were sub-metacentric.

Medini (*C. winterinaus*) is comparatively new variety than above two varieties of *C. winterianus* which is also classified under Java Citronella grass with 2*n*=20 as its somatic chromosome complement (Fig. 1e). Arm ratios for the chromosomes of this species ranged from 1.04 to 3.40 and the range of chromosome lengths lies between 1.70 to 3.10 µm (Fig. 2c). The length of two chromosome ranged between 2.0 to 3.0µm (2 C chromosomes) and other eight chromosomes were ranged in 1.0 to 2.0 µm (8 D chromosomes). As per the arm ratios of different chromosomes of this variety, 3 of them were characterized as metacentric whereas 7 of them were characterized as sub-metacentric (Table 1).

Krishna (*C. flexuosus*), also contains the somatic chromosome complement of 2*n*=20 (Fig. 1f). The arm ratios of the chromosomes of this species ranged from 1.07 to 2.22 and the range of chromosomes length lies between 1.48 to 2.30 µm (Figure 2d). This showed that *C. flexuosus* var. Krishna had slightly smaller chromosomes than *C. winterianus*. As per the arm ratios of different chromosomes of this variety, 5 of the chromosomes were metacentric and other 5 were sub-metacentric (Table 1).

### Localization of 45S rDNA

Bio-13 (*C. winterianus*) had shown strong hybridization signal of 45S rDNA at the terminal ends of short arms of the longest heterologous pair of somatic chromosome complement. The differential signal intensity of 45S rDNA was observed in different stages like metaphase (Fig. 1a and 1c) and interphase (Fig. 1b). The relative length of 45S rDNA hybridization signal in one chromosome of the heterologous bivalent was 29.48% (0.92µm) of the whole chromosome while the relative length of 45S rDNA in another chromosome of this pair was 10.01% (0.28 µm). Surprisingly, the signal intensity on both the chromosomes of this heterologous chromosome pair is not equal which is indicating the differential copy number of rDNA in both chromosomes of this heterologous chromosome pair.

Manjari (*C. winterianus*) had also shown strong hybridization signals of 45S rDNA at the terminal ends of short arms of longest heterologous pair of its chromosome complement (Fig 1d). Similar to the Bio-13, the differential signal intensity of 45S rDNA was observed in metaphase chromosomes of Manjari (Fig 1d). The relative length of 45S rDNA signal in one chromosome of the heterologous bivalent was 26.51% (0.79 µm), while the relative length of 45S rDNA in other chromosome of this pair was recorded 11.59 % (0.32 µm) which is indicating the differential copy number of rDNA in both chromosomes of this heterologous chromosome pair.

Medini (*C. winterianus*) had shown to have strong hybridization signals of 45S rDNA at the terminal ends of short arms of longest heterologous pair of its chromosome complement (Fig 1e). Similar to the Bio-13 and Manjari, this variety of *C. winterianus* also showed differential signal intensity of 45S rDNA in metaphase chromosomes. The relative length of 45S rDNA signal in one chromosome of the heterologous bivalent was 26.12% (0.81 µm) while the relative length of 45S rDNA in other chromosome of this pair was recorded 11.72% (0.32 µm) which is indicating the differential copy number of rDNA in both chromosomes of this heterologous chromosome pair. Krishna (*C. flexuosus*) had shown to have strong hybridization signals of 45S rDNA at the terminal ends of short arms of longest chromosome pair of its chromosome complement. However, in contrast to differential signal intensity of 45S rDNA shown in all the three varieties of *C. winterianus*, this species had shown to have the 45S rDNA signals of similar intensity i.e., 32.60% and 33.18% on both chromosomes of longest homologous chromosome pair (Fig 1f).

## Discussion

As reflected from the results, the arm ratios for the chromosome of *C. winterianus* ranged from 1.05 to 3.37 and the range of chromosomes length lies between 1.3 to 3.12 µm (Fig. 2). These results are in almost accordance of previous karyomorphological observation [14] where he reported the average arm ratio 1.26 and the chromosomal length in the range of 1.3 to 3.15 µm of the species *C. winterianus* Jowitt. As per the arm ratios of different chromosomes of all the genotypes of *C. winterianus*, 3 chromosomes are metacentric and 7 are sub-metacentric. These results are in contrast to the previous investigation [14] where *C. winterianus* was reported to have 8 sub-metacentric and 2 metacentric chromosomes. This difference may be due to small size of the chromosomes as sometimes it may be difficult to identify the centromeric position of the chromosomes. However, during the present investigation we have recorded the data on several cells and the data were based on average length and hence seems to be more authentic. The slight variation observed in the karyotypes of *C. winterianus* may be due to phenomenon of differential chromatin condensation in different chromosomes. Further, the lemon grass variety Krishna (*C. flexuosus*) has also shown its somatic complement as 2*n* = 20 with basic chromosome number *x =* 10 (Fig. 1f and 2d) which is very common in many of the species of *Cymbopogon*. The average chromosome arm ratios of *C. flexuosus* (1.95) are slightly varied from the arm ratio (1.53) of earlier reports [14].

In the past few decades, the extensive study of many plant species for localization of 45S rDNA genes have been done through FISH technique and it has been observed that, most of diploid plants have two sites rDNA i.e., a single locus [15]. Even though, some diploids exist with numerous sites of rDNA. The rDNA copy number changes rapidly and frequently, triggering relocation or deletion of some loci over and above the reduction in copy number below the detection sensitivity limit or mapping resolution [16-17]. For the justification of these differences some mechanisms have been proposed by various workers, such as various chromosome rearrangements, unequal crossing-over, gene transposition (gene mobility), and conversion [18-20]. There is certain proof that rDNA sites may alter chromosomal position, lacking the contribution of translocations [19]. Some of other observations have led to the suggestion that rDNA tandem repeats are mobile, with different molecular mechanisms believed to be intricated in this type of movement [21-22]. Signifying the transposition of the rDNA loci [23] triggered by En/Spm-like transposons [17,21,24], this procedure might designate chromosomal re-patterning of rDNA, which appears to be triggered through En/Spm-like transposons being accountable for drifting of rDNA to a new location [17]. Examples for arithmetical changes in 45S rDNA sites were described for few additional species, like in the colchicine-induced auto tetraploid *Arabidopsis thaliana* [25], signifying the 45S rDNA-bearing chromosome reorganization, as an entity of intragenomic translocation, and in tetraploid *Centaurea jacea* [26] in which the deletion of one pair of 45S rDNA loci was observed. Unluckily, there was lack of concrete evidence about mechanism involved in rDNA loci variation, so further studies need to be done in order to find the reasons behind the variation in pattern of rDNA loci.

The observations recorded during the present investigation are slightly different as the polymorphism has been observed within the same loci which exist at the short arms of two chromosomes of a homologous pair which seems like a rare observation, ‘as it is well known fact that in case of variation in rDNA present in two chromosomes of a homologous pair it will be equilibrated by subsequent round of crossing over and recombination in next generations. Krishna is a variety of *C. flexuosus* which is developed from variety Pragati and Cauveri (*C. flexuosus)* through phenotypic recurrent selection programme [27]. The crosses led to homogenization of genes through recombination and crossing over hence there is no polymorphism in rDNA sites (Fig. 1f and 2d) on both the homologous chromosomes and thus the intensity of rDNA hybridization signals remain similar in both chromosomes. However, Bio-13 (*C. winterianus*) was developed through mutation breeding. Manjari was developed by irradiation of the slips of variety Manjusha with gamma rays and Medini was developed by clonal selection of some of the well performing commercial varieties of *C. winterianus*. These facts prompted us to explore the possibilities for differential distribution of 45S rDNA in both chromosomes of a homologous chromosome pair existed in the above-mentioned varieties of *C. winterianus*.

Java citronella, i.e., *C. winterianus* flowers copiously in South India and at higher attitudes in the hills of North Eastern India. However, due to irregularities in meiosis and chromosome polyploidy, viable seeds are not formed and therefore, the species can be propagated only by vegetative means. As per perusal of different literature we come across the fact that heterochromatin sites are fragile in nature and are prone to degeneration upon the exposure of physical and chemical mutagens. Considering the importance of these species there are several programmes running in different institutions to increase quantity and quality aspect. The development of *C. winterianus* var. Manjari through gamma radiation may be responsible for breakage of fragile rDNA site from the chromosome or this variability may be due to evolutionary speciation due to natural cross among two different species of *Cymbopogon* which is fixed due to clonal propagation (Fig. 3). However, in both the situations these changes were fixed by vegetative method of propagation which is general mode of reproduction in the case of *C. winterianus*. Thus, the varieties of *C. winterianus* never got chance to recover by chromosome homogenization. This may be the reason why the change got fixed in cells of these varieties of *C. winterianus*. For the variety Medini of *C. winterianus* which is developed through clonal selection from Manjari and having the same background. The *C. winterianus* var. Bio-13 is developed by *in vitro* somaclonal selection [28]. However, seed setting in *C. flexuosus* var. Krishna takes place in all regions of India, and hence developed through cross pollination and recurrent selection. In addition to vegetative propagation, it is also being propagated through seeds, hence there is equal opportunity of chromosomal recovery in each generation.

**Figure 3.**
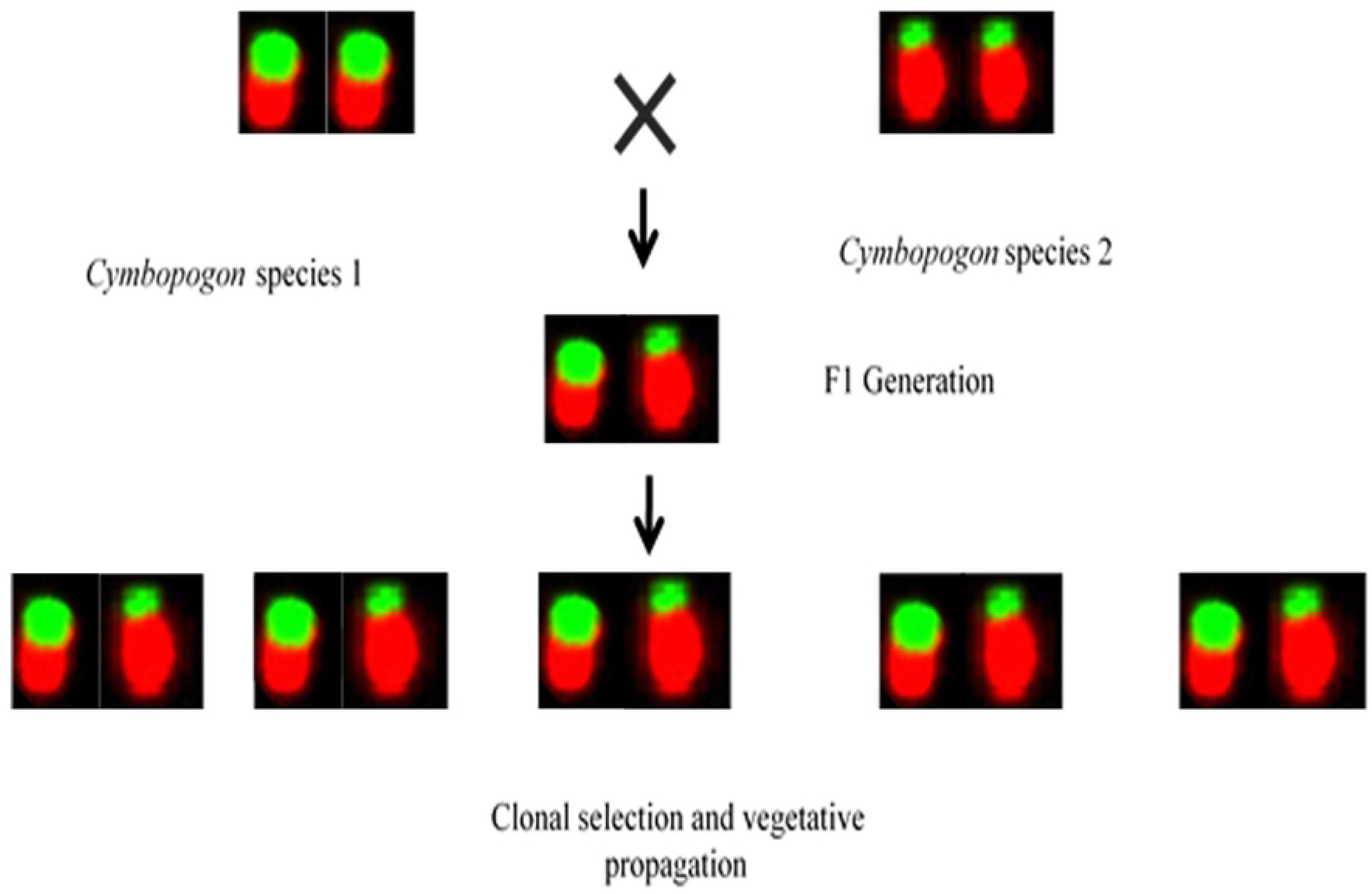
Evolutionary speciation in *C. winterianus*.

## Material and Methods

### Plant materials

Four varieties of two *Cymbopogon* species namely *C. winterianus* (Medini, Manjari, Bio-13) and *C. flexuosus* (Krishna) were collected from Central Institute of Medicinal and Aromatic Plants (CIMAP), Research Centre, Pantnagar, Uttarakhand, India. The information concerning the details of cultivars, chromosome number and percentage listed in Table 1.

### Preparation of chromosome spreads

The procedure of mitotic chromosome preparation was basically the same as published protocols [29] with some modifications. The slips of these cultivars were covered with moist paper and kept in a tray for root initiation in dark under room temperature for 68-72 hr. Lateral roots of about 1 cm were pre-treated in 0.002 mol/L 8-hydroxyquinoline at room temperature for 2 h, then fixed in 3:1 Carnoy’s fixative solution for at least 1 day. To obtain the chromosome preparations, fixed root tips were digested with enzyme mixtures, containing 4% cellulose Onozuka R-10 (Merck, http://www.merck-chemicals.com) and 2.0 % pectinase (Sigma-Aldrich, http://www.sigmaaldrich.com) in 1X PBS buffer, pH 5.5, at 37°C for 50 min. The enzyme solution was replaced by deionized water and kept on ice for at least 8 min. Following, the digested root tips were fixed in 3:1 Carnoy’s fixative. The slides were obtained using a “flame-dried” method, according to the published protocol [30].

### Probe preparation

Plasmid pTa71 with a size 9.0 kb from wheat [31] was used for 45S rDNA sites. rDNA was labelled with Fluorescein-12-dUTP (ROCHE Diagnostics) using the nick translation method.

### Signal detection and analysis

A 30 µl of hybridization mixture contained 15 µl of deionized formamide, 3 µl of 20X SSC, 6µl of 50% dextran sulphate, 1 µl of Salmon sperm DNA, 2 µl of probe DNA and 3 µl of ddH_2_O was added to denatured slide and covered with plastic coverslip. The slides were then incubated at 37° C in moist chamber overnight for hybridization of labelled probe with target DNA. After overnight incubation, slides were washed three times in 2X SSC for 5 min, 50% formamide in 2X SSC for 10 min and three times with 2X SSC for 5 min each at 42° C in water bath. This was followed by subsequent washing in 1X SSC for 5 min at RT. The slides were counterstained with propidium iodide and mounted with single drop of anti-fade mounting medium (Vectashield) and covered with 22×30 mm coverslips.

Photographs of cells were captured with well spread chromosomes by epifluorescence Zeiss Axioimager MI microscope (Carl Zeiss, Germany). The mean length of each chromosome, chromosome length, long arm, short arm and arm ratio of each chromosome and percentage of 45S rDNA signals were obtained through measurements with MicroMeasure 3.3 software (http://www.colostate.edu/Depts/Biology/MicroMeasure). The karyograms were developed using above parameters obtained through MicroMeasure 3.3.

## Acknowledgments

The author acknowledged the support of Central Institute of Medicinal and Aromatic Plants (CIMAP), Research Centre, Pantnagar, Uttarakhand, India for providing the germplasm. The research was performed at Molecular Cytogenetics Laboratory, Department of Molecular Biology & Genetic Engineering, College of Basic Sciences & Humanities, G. B. Pant University of Agriculture & Technology, Pantnagar, Uttarakhand.

---

**Table 2.**
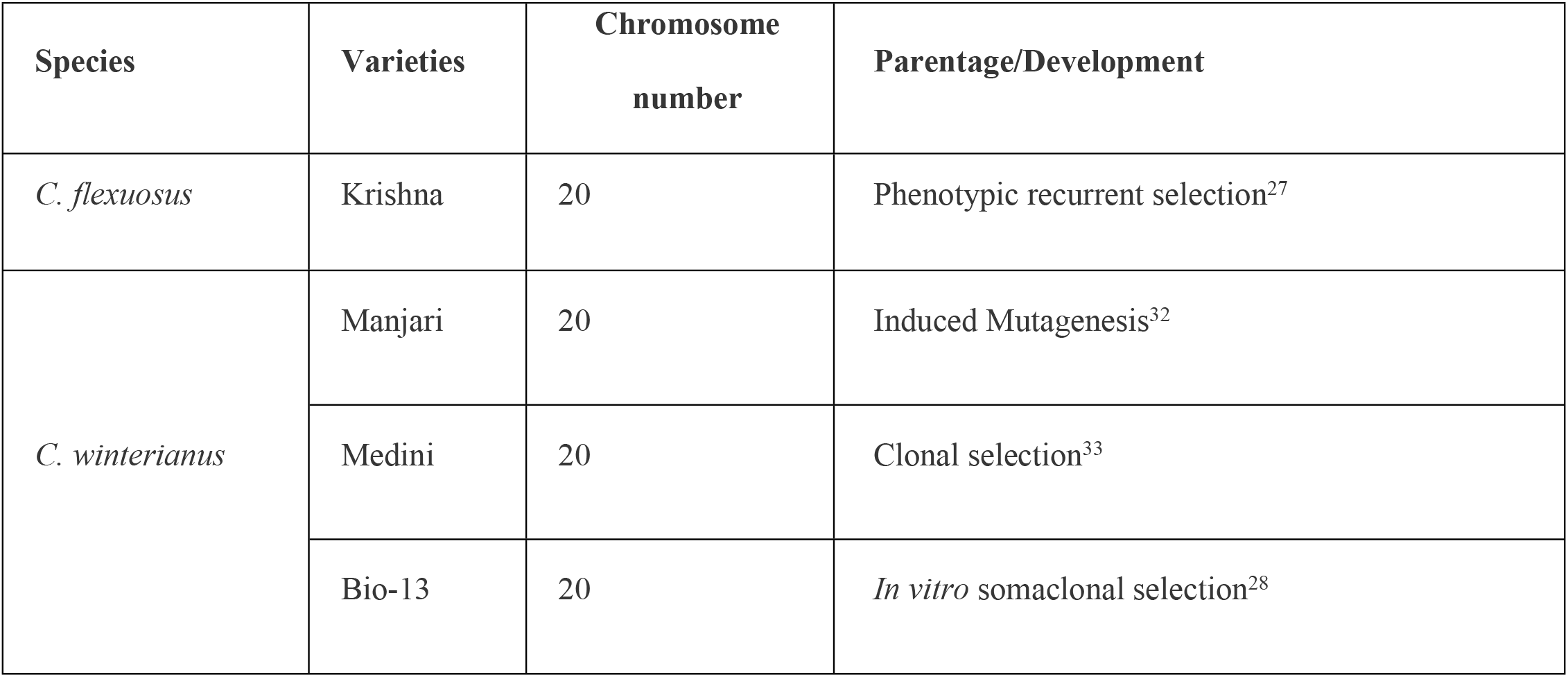
Chromosome number and parentage of *Cymbopogon* species used in the present study.

| Species | Varieties | Chromosome number | Parentage/Development |
| --- | --- | --- | --- |
| *C. flexuosus* | Krishna | 20 | Phenotypic recurrent selection<sup>27</sup> |
| *C. winterianus* | Manjari | 20 | Induced Mutagenesis<sup>32</sup> |
|  | Medini | 20 | Clonal selection<sup>33</sup> |
|  | Bio-13 | 20 | *In vitro* somaclonal selection<sup>28</sup> |

## References

1. Shasany A K, Lal RK, Darokar MP, Patra NK, Garg A, Kumar S, Khanuja SPS. Phenotypic and RAPD diversity among Cymbopogon winterianus Jowitt accessions in relation to Cymbopogon nardus Rendle. Genetic Resources and Crop Evolution 2000; 47(5): 553–9. https://doi.org/10.1023/A:1008712604390.

2. Rao BL. Scope for development of new cultivars of Cymbopogon as a source of terpene chemicals. In: Handa SS, Kaul MK, editors. Supplement to cultivation and utilization of aromatic plants. National Institute of Science Communication, Dr. KS. Krishnan Marg, New Delhi, India; 1997.p. 71–83.

3. Babu CN. Chromosome numbers in Cymbopogon species. Current Science 1936; 4(10): 739–40.

4. Bor NR. The genus Cymbopogon Spreng. In India Burma and Ceylon Part I. Journal of the Bombay Natural History Society 1953; 51: 890–916.

5. Sangwan NS, Yadav U, Sangwan RS. Molecular analysis of genetic diversity in elite Indian cultivars of essential oil trade types of aromatic grasses (Cymbopogon species). Plant Cell Reports 2001; 20(5): 437–44. https://doi.org/10.1007/s002990100324.

6. Kumar J, Verma V, Goyal A, Shahil AK, Sparoo R, Sangwan RS, Qazi GN. Genetic diversity analysis in Cymbopogon species using DNA markers. Plant Omics Journal 2009; 2(1): 20–9.

7. Mukai Y, Endo TR, Gill BS. Physical mapping of the 18S-26S rRNA multigene family in common wheat: identification of a new locus. Heredity 1991; 100(100):71–8 https://doi.org/10.1007/bf00418239.

8. Leitch IJ, Heslop-Harison, JS. Physical mapping of the 18S– 5.8S–26S rDNA genes in barley by in situ hybridization. Genome 1992; 35(6): 1013–18. https://doi.org/10.1139/g92-155.

9. Pedersen C, Linde-Laursen I. Chromosome location of four minor rDNA loci and marker microsatellite sequence in barley. Chromosome 1994; 2(1): 65–71. https://doi.org/10.1007/bf01539456.

10. Badaeva ED, Bernd F, Gill BS. Genome differentiation in Aegilops. Physical mapping of 5S and 18S-26S ribosomal RNA gene families in diploid species. Genome 1996; 39(6): 1150–58 https://doi.org/10.1139/g96-145.

11. Raina SN, Mukai Y. Detection of a variable number of 18S–5.8S–26S and 5S ribosomal DNA loci by fluorescence in situ hybridization in diploid and tetraploid Arachis species. Genome 1999; 42(1): 52–59. https://doi.org/10.1139/g98-092.

12. Ansari HA, Ellison NW, Reader MS, Badaeva ED, Friebe B, Miller TE, Williams WM. Molecular Cytogenetic Organization of 5S and 18S-26S rDNA Loci in White Clover (Trifolium repens L.) and Related Species. Annals of Botany 1999; 83(3): 199–206 https://doi.org/10.1006/anbo.1998.0806.

13. Taketa S, Ando H, Takeda K, Bothmer VR. Physical locations of 5S and 18S–25S rDNA in Asian and American diploid Hordeum species with the I genome. Heredity 2001; 86(5): 522–30 https://doi.org/10.1046/j.1365-2540.2001.00768.x.

14. Lavania UC. Karyomorphological observations in Cymbopogon Sprengel. Cytologia 1988; 53(3): 517–24. https://doi.org/10.1508/cytologia.53.517.

15. Mishima M, Ohmido N, Fukui K, Yahara T. Trends in site number change of rDNA loci during polyploid evolution in Sanguisorba (Rosaceae). Chromosoma 2002; 110(8): 550 –8. https://doi.org/10.1007/s00412-001-0175-z.

16. Jiang J, Gill BS. New 18S.26S ribosomal RNA gene loci: chromosomal landmarks for the evolution of polyploidy wheats. Chromosoma 1994; 103(103):179–185. https://doi.org/10.1007/s004120050022.

17. Raskina O, Belyayev, A. and Nevo, E. Activity of the En/Spm-like transposons in meiosis as a base for chromosome re-patterning in a small, isolated, peripheral population of Aegilops speltoides Tausch. Chromosome Research 2004; 12(2): 153–61.

18. Schubert I, Wobus U. In situ hybridization confirms jumping nucleolus organizing regions in Allium. Chromosoma 1985; 92(2): 143–48. https://doi.org/10.1007/bf00328466.

19. Dubcovsky J, Dvorak J. Ribosomal RNA multigene loci: nomads of the Triticeae genomes. Genetics 1995; 140(4): 1367–77.

20. Thomas HM, Harper JA, Morgan WG. Gross chromosome rearrangements are occurring in an accession of the grass Lolium rigidum. Chromosome Research 2001; 9 (7): 585–90 https://doi.org/10.1023/A:1012499303514.

21. Datson PM, Murray BG. Ribosomal DNA locus evolution in Nemesia: transposition rather than structural rearrangement as the key mechanism. Chromosome Research 2006; 14(8): 845–857. https://doi.org/10.1007/s10577-006-1092-z.

22. Pedrosa-Harand A, De Almeida CCS, Mosiolek M, Blair MW, Schweizer D, Guerra M. Extensive ribosomal DNA amplification during Andean common bean (Phaseolus vulgaris L.) evolution. Theoretical and Applied Genetics 2006; 112 (5): 924–33. https://doi.org/10.1007/s00122-005-0196-8.

23. Frello S, Heslop-Harrison JS. Chromosomal variation in Crocus vernus Hill (Iridaceae) investigated by in situ hybridization of rDNA and a tandemly repeated sequence. Annals of Botany2000; 86(2): 317–22. https://doi.org/10.1006/anbo.2000.1189.

24. Raskina O, Belyayev A, Nevo E. Quantum speciation in Aegilops: Molecular cytogenetic evidence from rDNA cluster variability in natural populations. Proceedings of National Academy of Sciences of the USA 2004; 101(41): 14818–14823.

25. Weiss H, Maluszynska J. Chromosomal rearrangement in autotetraploid plants of Arabidopsis thaliana. Hereditas 2000; 133(3): 255–61. https://doi.org/10.1111/j.1601-5223.2000.00255.x.

26. Dydak M, Kolano B, Nowak T, Siwinska D, Maluszynska J. Cytogenetic studies of three European species of Centaurea L. (Asteraceae). Hereditas 2009; 146 (4): 152–61. https://doi.org/10.1111/j.1601-5223.2009.02113.x.

27. Improved Clonal variety Krishna of lemongrass developed. CIMAP Newsletter 1997; 24: 2–3.

28. Patra NK, Kumar B. Improved Varieties and Genetic Research in Medicinal and Aromatic Plants (MAPs). In: Proceedings of Second National Interactive Meet (NIM) on Medicinal and Aromatic Plants, Lucknow, India 2005; 53–61.

29. Lou QF, He YH, Cheng CY, Zhang ZH, Li J, Huang, SW Chen JF. Integration of highresolution physical and genetic map reveals differential recombination frequency between chromosomes and the genome assembling quality in cucumber. Plos ONE 2013; 8(5): e62676. doi: 10.1371/journal.pone.0062676. PMID: 23671621.

30. Iovene M, Wielgus SM, Simon PW, Buell CR, Jiang JM. Chromatin structure and physical mapping of chromosome 6 of potato and comparative analysis with tomato. Genetics 2008; 180(3): 1307–17.

31. Gerlach WN, Bedbrook JR. Cloning and characterization of ribosomal RNA genes from wheat and barley. Nucleic Acids Res 1979; 7(7): 1869–85.https://doi.org/10.1093/nar/7.7.1869.

32. Lal RK, Sharma JR, Misra HO. Development of new variety Manjari of Citronella Java (C. winterianus). Journal of Medicinal and Aromatic Plant Science 1999; 21: 727–29.

33. High yielding varieties Jalpallavi, Manjari and Medini of Citronella Java (Cymbopogon winterianus), CIMAP Newsletter 1999; 26: 3–5.

